# Song type variations of Louisiana Waterthrush (*Parkesia motacilla*) and their geographic distributions

**DOI:** 10.1101/2020.09.24.312454

**Authors:** W. Ross Silcock, Shari L. Schwartz, John U. Carlini, Stephen J. Dinsmore

## Abstract

Louisiana Waterthrush *(Parkesia motacilla)* is a familiar singer in the Western Hemisphere family *Parulidae*, yet apparent geographic variations in its song and potentially related causal mechanisms have not received detailed examination in previously published studies. Here, we analyzed song pattern variations of 651 Louisiana Waterthrush singers in audio spectrogram recordings obtained from our field work and publicly accessible bioacoustics archives. Visual and auditory assessment of the introductory note sequence of each song identified three distinct song types (A, B, and C) and 88.3% of the songs were assigned to one of these types. Linear Discriminant Analysis and Random Forest methods were used to verify the assignments and showed strong agreement (>90%) for Type A with slightly less agreement on Types B and C. User error rates (proportion of the Linear Discriminant Analysis classifications that were incorrect) were <10% for Types A and B, but 26% for Type C, while producer error rates (proportion of the song type for which the Linear Discriminant Analysis was incorrect) were >25% for Types A and C, but <5% for Type B. Our findings confirmed in a subset of 87 individuals that most between-individual variation was in the number of notes and note sequence duration while most within-individual variation resulted from the percent of downstrokes. The location of each singer was plotted on a map of the breeding range and results indicated the song types have large-scale discrete geographic distributions that co-occur in some regions but not range-wide. Evaluation of the distributions provided tentative support for a hypothesis that two of the song types may independently exhibit congruence with the geographic extent of Pleistocene glacial boundaries and the third song type may be distinguished by a lack of congruence, but further investigation is needed to elucidate whether the song variations represent subpopulations with three separate evolutionary histories.

## Introduction

Geographic variation in songbird vocalizations has been documented in nearly all rigorously studied species [1, 2] and can be the consequence of various factors [3]. Imitative learning promotes geographic variation when cultural transmission of songs between generations allows shared novel vocalization patterns in song structure to arise through inaccurate learning and appearance of new components [4––6]. Nevertheless, variation in vocalizations is not unrestrained. Instead of copying the range of sounds to which they might be exposed, juvenile oscines preferentially learn the vocalizations characteristic of their species [7] or subspecies [8], and variations that arise during song evolution may be constrained or redirected by genetically-based mechanisms such as physiological song production abilities [3]. Variation may also be limited within small colonizing or isolated populations that arise from genetically depauperate founder populations such as those on islands or in recently deglaciated areas populated from glacial refugia [9—12]. Populations that become geographically isolated undergo random cultural drift [13], and the songs of isolates may ultimately evolve to the extent that conspecifics in other populations alter their responses which can lead to reproductive isolation [14] and divergence as was found with Australia’s Chowchilla (*Orthonyx spaldingii*) whose song structure has large-scale geographic variations in bandwidth and frequency peaks attributed to cultural drift during isolation in Pleistocene refugia [15].

A combination of multiple evolutionary processes may influence geographic song variation as was observed in the bandwidth and internote duration variances between eastern and western Common Yellowthroats (*Geothlypis trichas*) by Bolus [16], who noted sexual selection can be a contributor that reinforces changes in song structure. Although nearly a third of all female oscines are not known to sing [17], they must nonetheless learn their parental song type alongside their male siblings in order to recognize and select a conspecific mate [18, 19]. Song dialects, defined by Mundinger [20] as variation between song forms with loosely discrete boundaries, can be distinguished by both males and females of White-crowned Sparrow (*Zonotrichia leucophrys*) subspecies *nuttalli* [21]. Kroodsma [22] suggested certain populations with songs that differ discontinuously may possess differing evolutionary histories as has been demonstrated by the closely related and phenotypically similar taxonomic pairs of Marsh Wrens (*Cistothorus palustris plesius and C. p. iliacus*) [22, 23] and Eastern and Western meadowlarks (*Sturnella magna* and *S. neglecta*) [24, 25]. In regions of recontact, singing eastern and western Marsh Wrens and Eastern and Western meadowlarks evoke territorial defense from both respective male congeners despite their contrasting songs, yet females of both pairs of taxa reserve receptivity responses solely for singers of their parental song type [22, 24]. The importance of female discrimination was demonstrated in a study of sympatric Eastern and Western meadowlarks [26] in which the absence of song convergence was attributed to selection in order to oppose the establishment of a single song type and avoid interspecific matings in the area of sympatry.

Song divergence has been correlated with geographic variation in numerous avian subpopulations [16, 27—29]. It has been assumed that Louisiana Waterthrush exhibits no phenotypic or genotypic geographic variation [30] in contrast with sister species Northern Waterthrush (*Parkesia noveboracensis*) [31], although Mattsson et al. [30] proposed subpopulations could potentially arise from discontinuous waterways on breeding grounds. A lack of connectivity between some riparian corridors may limit dispersal as suggested by Mattsson [32] for Louisiana Waterthrush, and by Machtans for other forest songbirds [33]. Geographic discontinuities in song structure can occur in species that have a pattern of limited dispersal and learn relatively simple repertoires in the natal region during the pre-dispersal period [3], the latter characteristic having been documented in Louisiana Waterthrush [30, 34, 35].

Studies have revealed that artifactual adaptations to Pleistocene glaciations still endure today in some Neotropical migrants as exemplified by the unnecessarily circuitous migratory pathway that the Swainson’s Thrush (*Catharus ustulatus*) has retained [36]. The ~2.6 million year Pleistocene Epoch [37] not only affected heritable traits, but also determined the contemporary distributions of many avian species [38—40] as repeated cycles of advancing and retreating glacial ice [41, 42] fragmented and redistributed vegetation types [43] such as on the Great Plains where habitat transitioned between forests and grasslands [44]. Range expansions and recolonizations across newly-deglaciated habitat during ~20,000-year interglacial periods were succeeded by glacial intrusions that separated and shifted populations into discrete refugia for ~100,000-year periods [45]. East of the Rocky Mountains across what is presently Louisiana Waterthrush’s breeding range [46], nearly all of the Laurentide Ice Sheet reached maximum extent during the earlier pre-Illinoian Stage glaciations, whereas only the central and eastern portions of the ice sheet reached commensurately extensive glacial maxima during the later Illinoian Stage, and only the eastern portion of the ice sheet reached a comparable extent during the Last Glacial Maximum (LGM) of the most recent Wisconsinan Stage [42, 47]. During major glaciations, high-altitude periglacial permafrost in the Appalachian Mountains [48, 49] would have adversely affected habitability adjacent to the ice sheet. Periglacial permafrost conditions at lower elevations directly west of the Appalachians were restricted to a narrow band which allowed temperate forest vegetation to exist close to the ice margin, but farther west, permafrost conditions along the western edge of the ice sheet covered a wider expanse during the LGM [43, 50, 51]. Glacial vicariance impacted many aquatic and terrestrial taxa in the southeastern United States [52—55] during periods of glacial maxima as the ice sheet at maximum extent adjoined the Mississippi Embayment alluvial landscape [56, 57] and divided southeastern refugia into eastern and western locations referred to as Appalachian and Ozark centers of endemism by Strange and Burr [58]. The isolating combination of ice sheet and Embayment’s habitat barrier produced divergent lineages of North American salamanders [59, 60] and stream dwelling “highland fishes” [58, 61], all of which share aquatic invertebrate prey sources with Louisiana Waterthrush and may themselves be included in its diet [30]. The Embayment’s bisection of the distributions of some *Parulidae* species is evident in the current breeding ranges of Louisiana Waterthrush, Prairie Warbler (*Setophaga discolor*), and Worm-eating Warbler (*Helmitheros vermivorum*) [46].

As a result of observing Louisiana Waterthrush song pattern variations locally, we accessed publicly available bioacoustic archival recordings to inventory audio spectrograms from across the species’ entire breeding range. In this study, we analyzed the song types identified in the recordings, plotted their geographic distributions, assessed conceivable explanations for the configuration of the distributions, and evaluated each song type’s distribution for potential correlation with Pleistocene glacial conditions.

## Materials and methods

### Louisiana Waterthrush ecology

The Louisiana Waterthrush is unique among *Parulidae* as the only bird in the southeastern United States that breeds exclusively along forested streams [32]; prime habitat consists of narrow linear corridor territories along uncontaminated headwater streams in unfragmented mature forest with open understory and an optimal percentage of 30-69% deciduous species [62]. Both males and females have been observed returning to their same territory in consecutive years [30]. Most foraging time (88.0-90.4%) is spent in or beside the stream, and singing may be delivered from a favored perch above the stream or from the ground while foraging, and at times also in flight during territorial disputes [30]. The extent to which males discriminate between “familiar and unfamiliar” songs is not known [30]. Males sing vigorously upon arrival on the breeding grounds but after females arrive and are paired with males, singing sharply declines [34, 63]. Eaton [34] reported rarely hearing the “advertising song” from the time of pair formation until incubation started around 3-4 weeks later [30], with reinstated song never delivered as often as prenuptial singing. Males sing infrequently during the incubation and early nestling phases, and nestlings may on occasion also be exposed to female song [30]. Several females have been observed singing a version of male song that is recognizable but has a less complicated and softer delivery and apparently functions to summon the male during incubation recess [30]. Males at the latitude of Ithaca, New York sang occasionally in late June and early July while tending semi-independent juveniles, and had a resurgence of song in the first half of August after their molt was complete just prior to their early departure for wintering grounds [34]. The dependency period for fledglings lasts 3-4 weeks [30] after which some but not all may begin to wander from the natal area to varying degrees. Two juveniles banded by Eaton [34], were documented 2.0 km and 4.8 km from their natal areas just over a month after leaving the nest.

### Song structure

The repertoire of Louisiana Waterthrush is limited to a single song which is deemed to be its primary song when delivered without the extended conclusion that is appended occasionally [30]. Each male’s song is unique and individually recognizable [63, 64]. Variation in the pattern of introductory notes sung by some individuals has been noted [35]. Typical primary songs commence with a repetitive series of introductory notes and culminate in a phrase of non-repetitive notes [35]. The extended song conclusion can be appended to the primary song [65] singly or in repetition and is usually included when the singer is in territorial defense mode [35]. It is thought that when this extended song is added to the primary song, Louisiana Waterthrush’s song performs the same functions that some *Parulidae* species achieve by using a repertoire of more than one song [65, 66]. Unpaired Louisiana Waterthrush males often conclude the primary song without the extended song [64] while paired males appear to be more apt to add the extended conclusion [30, 65].

### Song type characterization

To investigate the range-wide distributions of the song type variations we had encountered in field work, all Louisiana Waterthrush audio recordings at Macaulay Library [67] and xeno-canto [68] from 1957 through 2018 were inventoried, and 820 spectrograms containing 4000+ songs were obtained. Visual and auditory examination of note shapes and sequences in the spectrograms allowed for identification of recordings shared by counter singing individuals whose songs required separate evaluation, and multiple recordings of a single singer submitted from the same location. After the elimination of recordings that were redundant, those that lacked sufficient quality for measurement, and a few from wintering grounds south of the United States, the resulting database (S1 Appendix) contained 3833 songs of 651 individual singers. For spectrograms containing more than one song, the RANDBETWEEN function in Microsoft Excel was used to select one song per singer at random. Only the series of introductory notes (“note” defined as a continuous trace on a spectrogram) that initiates the primary song was evaluated; we refer to this series as the Introductory Note Sequence (INS) (Fig 1). Raven Lite 2.0 sound analysis software (Cornell Laboratory of Ornithology) was utilized to measure the INS in each of the 651 songs for the seven following variables: number of notes (“Notes”), note sequence duration (“Duration”), duration per note (“Dur_note”), percent of duration spent singing the downstroke portions of notes (Percent_down”), minimum frequency (Min_freq”), maximum frequency (“Max_freq”), and frequency span (“Freq_span”). The phrase of notes that follows the INS and terminates the primary song is referred to as the Secondary Note Sequence (SNS); this phrase was distinguished from the INS by its miscellany of notes that extend to lower minimum frequencies than the minimum frequencies of the more uniformly patterned INS notes. SNS patterns shared regionally by multiple singers provided an additional aid for distinguishing the SNS from the INS, but because these apparent SNS geographic distributions plotted independently of INS distributions, that part of the primary song was not analyzed. The extended song conclusion that is appended inconsistently to the primary song was also excluded from analysis. Some singers uttered either a single call note or a brief “grace note” preceding the INS delivery but many singers did not employ these notes that often were too abbreviated to yield tonality. Some individuals sang INS notes with faint polyphonic signatures stacked vertically above the notes in the spectrogram image but these marks were presumed to indicate overtone frequencies so were not evaluated. After careful visual and auditory examination, the note shapes in the INS of each song were assigned to one of the three song types, or in cases where assignment was unclear, categorized as Unassigned.

**Fig 1.**
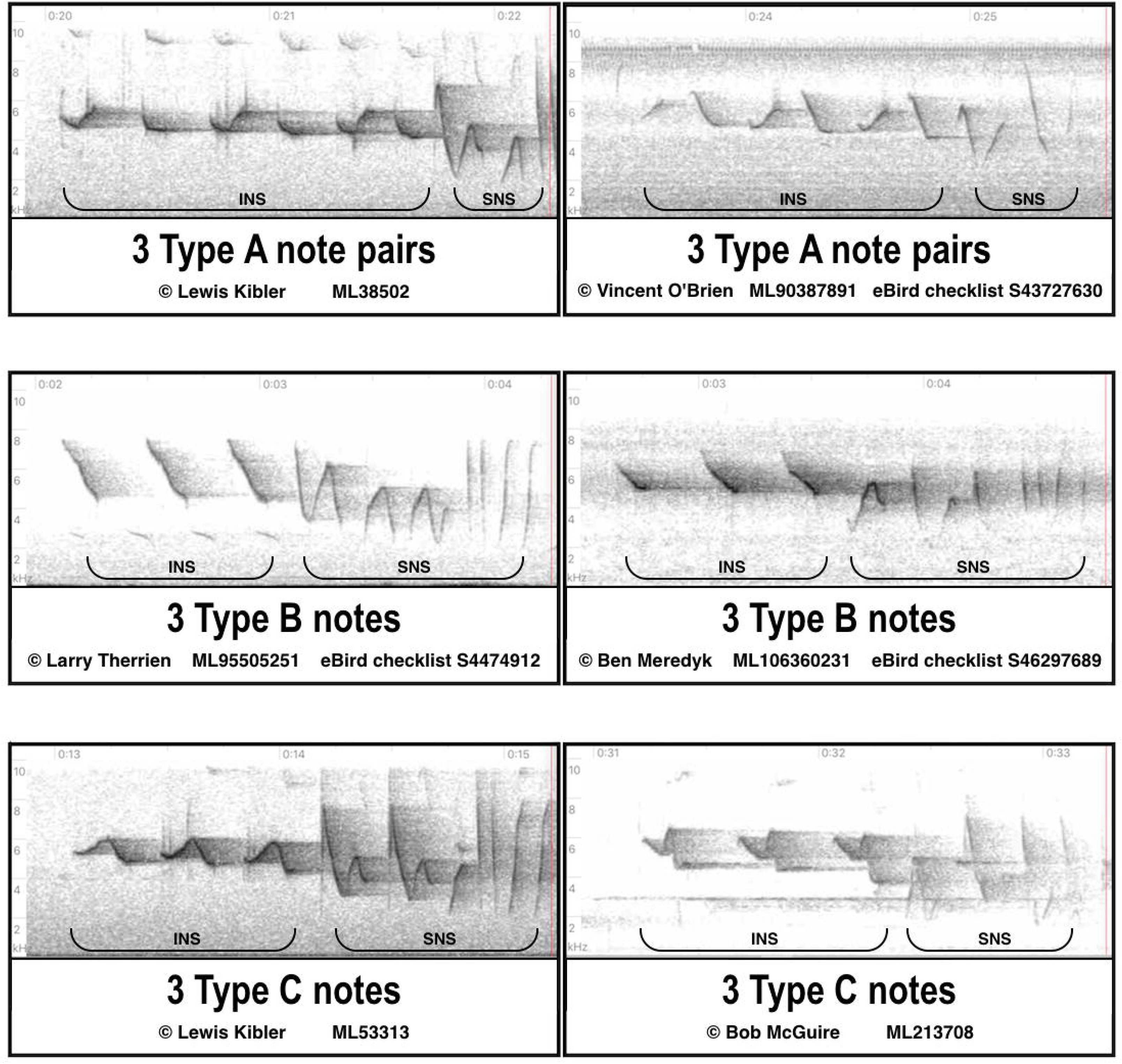
Spectrogram examples of the A, B, and C note types. Two examples (from S1 Appendix) are shown of each note type. Type A is comprised of alternating upstroke and downstroke note pairs, Type B employs downstrokes only, and Type C notes undulate both upwards and downwards.

### Statistical Analyses

Our goal was to identify and describe key characteristics of Louisiana Waterthrush songs, confirm the accuracy of our subjective classification of three song types, and understand which (if any) of seven song features were good predictors of song type. We used descriptive statistics (mean + SD) to characterize seven quantitative measures of song type. We then used Linear Discriminant Analysis (LDA) [69] to classify song type on the basis of the seven measures. For this analysis, we eliminated frequency span because it was strongly correlated with measures of minimum and maximum frequency, was highly variable, and was generally uninformative with respect to song type. Because the six remaining variables had different scales, we standardized them (SD = 1 for all of them) and then plotted them for visual inspection. This plot helped us interpret the influence of each of the six predictors of song type on a two-dimensional plot of song types for this analysis. For example, observations with a high value of Notes (above the mean) and a low value of Duration (below the mean) will be plotted in the bottom left of the LDA scores plot (e.g., in the direction of Notes and opposite the direction of Duration). Similarly, variables like Max_freq and Min_freq that are close to the origin (0,0), even after rescaling, have little impact on the position of the observations in the LDA scores plot. This was also confirmed with the use of a Random Forest (RF) analysis [70], which is similar to the LDA but allows interactions between measures. Lastly, we used multiple song measures to describe the intra- and inter-song variation in a sample of Louisiana Waterthrush spectrograms. We fit a variance-covariance model to the data where individual was a random variable and there were replicates within each individual. We used the Intra-Class Correlation (ICC), which is the group variance divided by total variance, to assess the proportion of variation that was due to within- and between-individual variation.

### Map of assigned singer locations with glacial environment

Recording locations were represented on a map of the eastern United States on Google My Maps and National Institute of Statistics and Geography map data (https://www.google.com/maps/d/u/0/viewer?mid=1PAkxNYqk_7iRslxEd57mgqR38u0&ll=40.669561163969384%2C-83.80830025&z=5) with color-coded pin markers placed at described locales or latitude and longitude coordinates provided by recordists. The 11 singers recorded in Florida are presumed migrants but were retained to contribute additional songs for analysis. Pleistocene glacial maximum boundaries were indicated on the map with gray screens that depicted the pre-Illinoian and Illinoian major glacial stages and the Wisconsinan LGM stage was represented with a thick black outline; map sources [42, 47, 71—75] encompassed both general and detailed illustrations to most accurately delineate former glacial boundaries for comparison to contemporary Louisiana Waterthrush locations. The high-altitude periglacial permafrost region in the Appalachian Mountains adjacent to the glacial boundary [48] was also depicted with a gray screen, the Wisconsinan LGM periglacial extent was depicted with a white line [50, 51], and the Mississippi Embayment region [74] that bisected suitable habitat in tandem with the ice sheet was portrayed with a thin black outline.

## Results

We measured 3833 Louisiana Waterthrush songs representing 651 individual singers (S1 Appendix) in this study. Visual/auditory assignments placed 88.33% of the songs into one of three song type groups designated as Type A (72; 11.06%), Type B (404; 62.06%), or Type C (99; 15.21%). The remaining 11.67% were categorized as Unassigned (“Un”: Table 1) and were omitted from the analyses depicted below in Figs 2, 4, and 5 and Tables 2 and 3.

**Table 1.** Summary statistics for measures of Louisiana Waterthrush song types.

| Type | n | Notes | Duration | Dur_note | Pct_down | Min_freq | Max_freq | Freq_span |
| --- | --- | --- | --- | --- | --- | --- | --- | --- |
| A | 72 | 4.40<br>(1.27) | 1.03<br>(0.25) | 0.24<br>(0.05) | 30 (9) | 4019<br>(273) | 6056<br>(481) | 2037<br>(478) |
| B | 404 | 2.81<br>(0.58) | 0.77<br>(0.20) | 0.27<br>(0.04) | 53 (9) | 4442<br>(230) | 6853<br>(455) | 2411<br>(441) |
| C | 99 | 2.86<br>(0.61) | 0.84<br>(0.20) | 0.30<br>(0.05) | 33 (9) | 4248<br>(259) | 6373<br>(451) | 2124<br>(465) |
| Un | 76 | 3.67<br>(1.28) | 0.94<br>(0.25) | 0.27<br>(0.05) | 37 (11) | 4187<br>(423) | 6443<br>(583) | 2256<br>(571) |

Numbers are means (SD) and are given for the seven quantitative measures of each song.

**Table 2.**
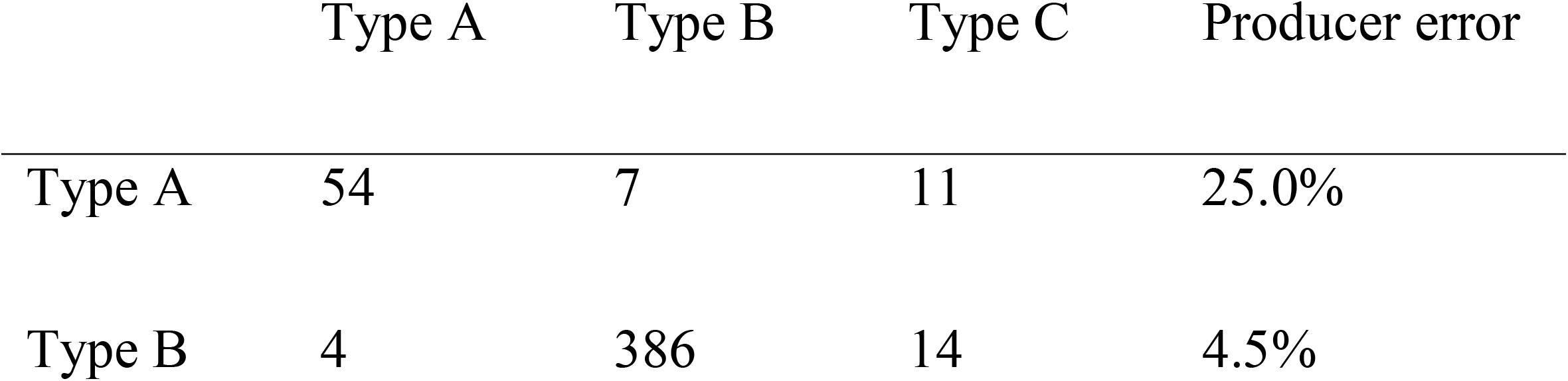
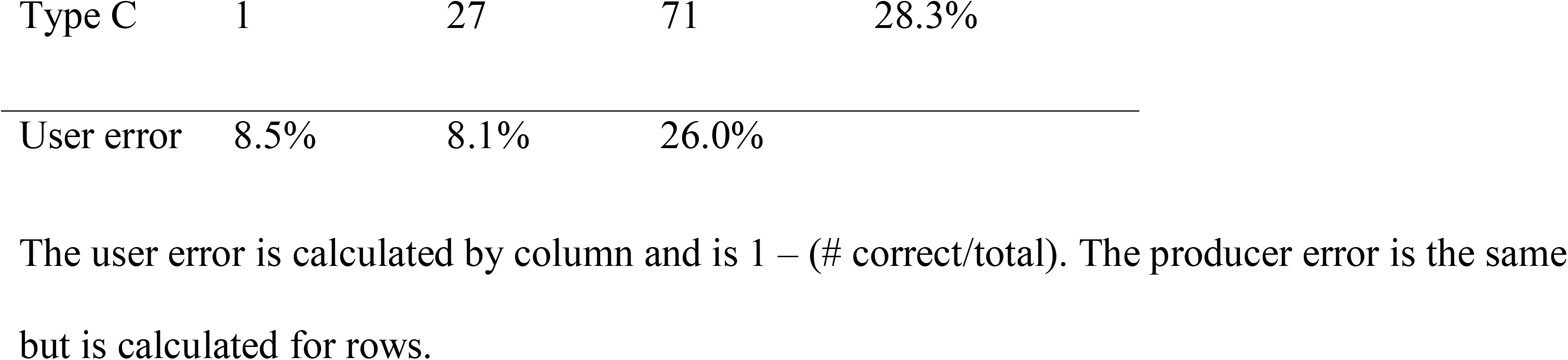
Confusion matrix for Louisiana Waterthrush song types from a Linear Discriminant Analysis of seven song attributes.

**Table 3.** Comparison of results of Linear Discriminant and Random Forest analyses for Louisiana Waterthrush song types.

| Style | Type A |  |  | Type B |  |  | Type C |  |  |
| --- | --- | --- | --- | --- | --- | --- | --- | --- | --- |
|  | A | B | C | A | B | C | A | B | C |
| Type A | 52 | 1 | 1 | 2 | 1 | 1 | 1 | 0 | 0 |
| Type B | 0 | 6 | 1 | 0 | 381 | 5 | 0 | 23 | 4 |
| Type C | 4 | 1 | 6 | 0 | 4 | 10 | 4 | 6 | 61 |

Rows are the Linear Discriminant Analysis assignments and columns are the Random Forest assignments.

**Fig 2.**
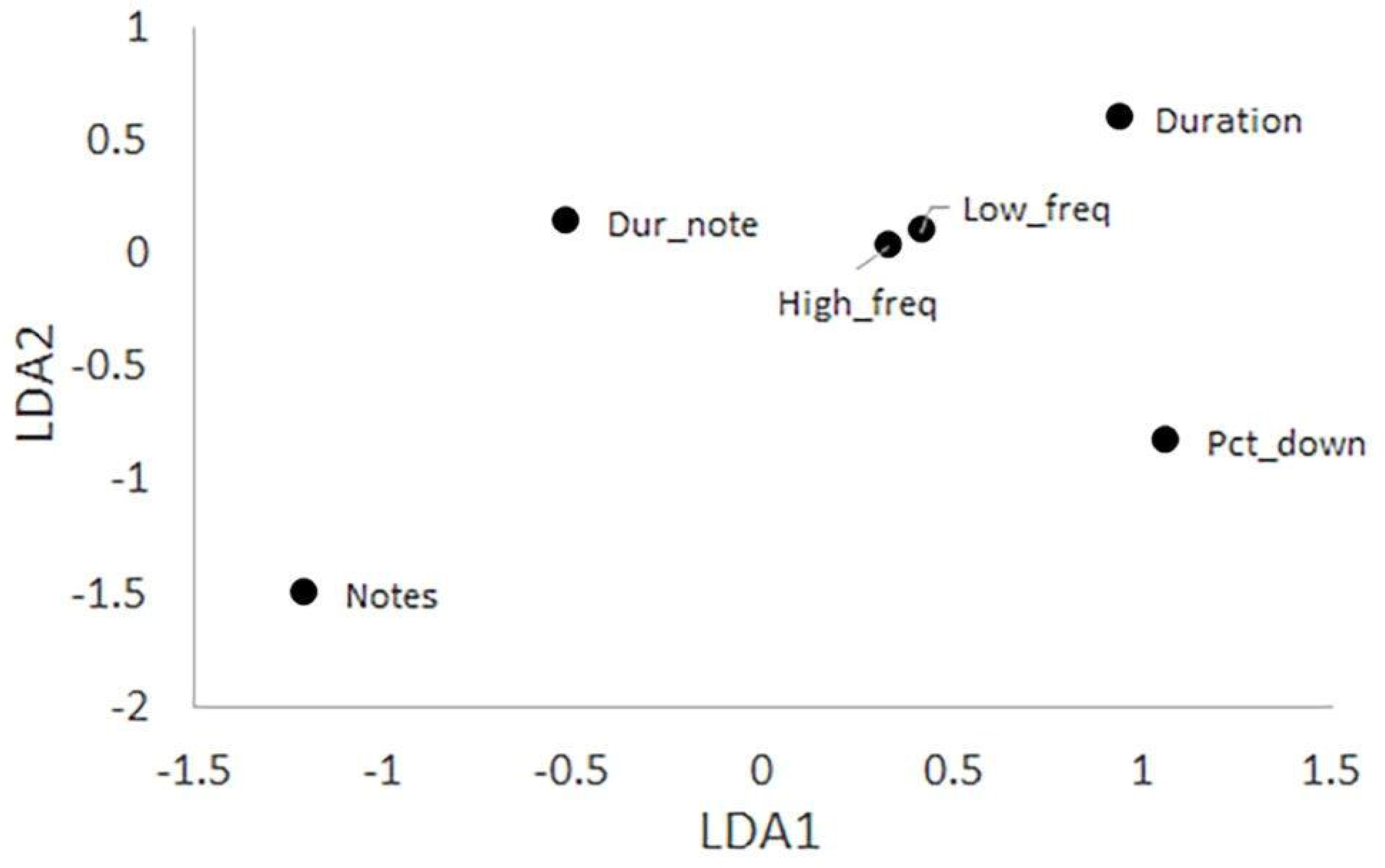
Plot showing the scaling coefficients of the Linear Discriminant Analysis of Louisiana Waterthrush song types. The six predictors of song type are arranged around the overall scaled mean (0,0) to indicate their relative influences. The six variables plotted are the number of notes (Notes), note sequence duration (Duration), the duration per note (Dur_note), the percent of duration spent singing the downstroke portions of notes (Pct_down), the minimum frequency (Min_freq), and the maximum frequency (Max_freq).

### Song type classification

For this analysis we used a subset of 651 songs, one per individual, for which we measured seven attributes (Table 1). The LDA coefficients varied across the six included variables (Fig 2) and helped interpret a plot of song type on the two LDA axes (Fig 3). Type A songs have above average values for Notes and below average values for Duration; the other four variables had little role in pulling Type A songs away from the other song types (Fig 3). The confusion matrix for three song types (Table 2) showed that the model performed well with most misclassifications occurring between Types B and C. The user error rates (proportion of the LDA classifications that were incorrect) were <10% for Types A and B, but 26% for Type C. The producer error rates (proportion of the song type for which the LDA was incorrect) were >25% for Types A and C, but were <5% for Type B. The LDA equations were:

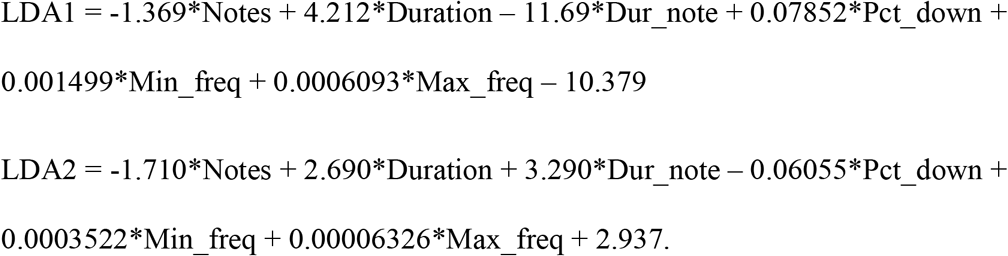

**Fig 3.**
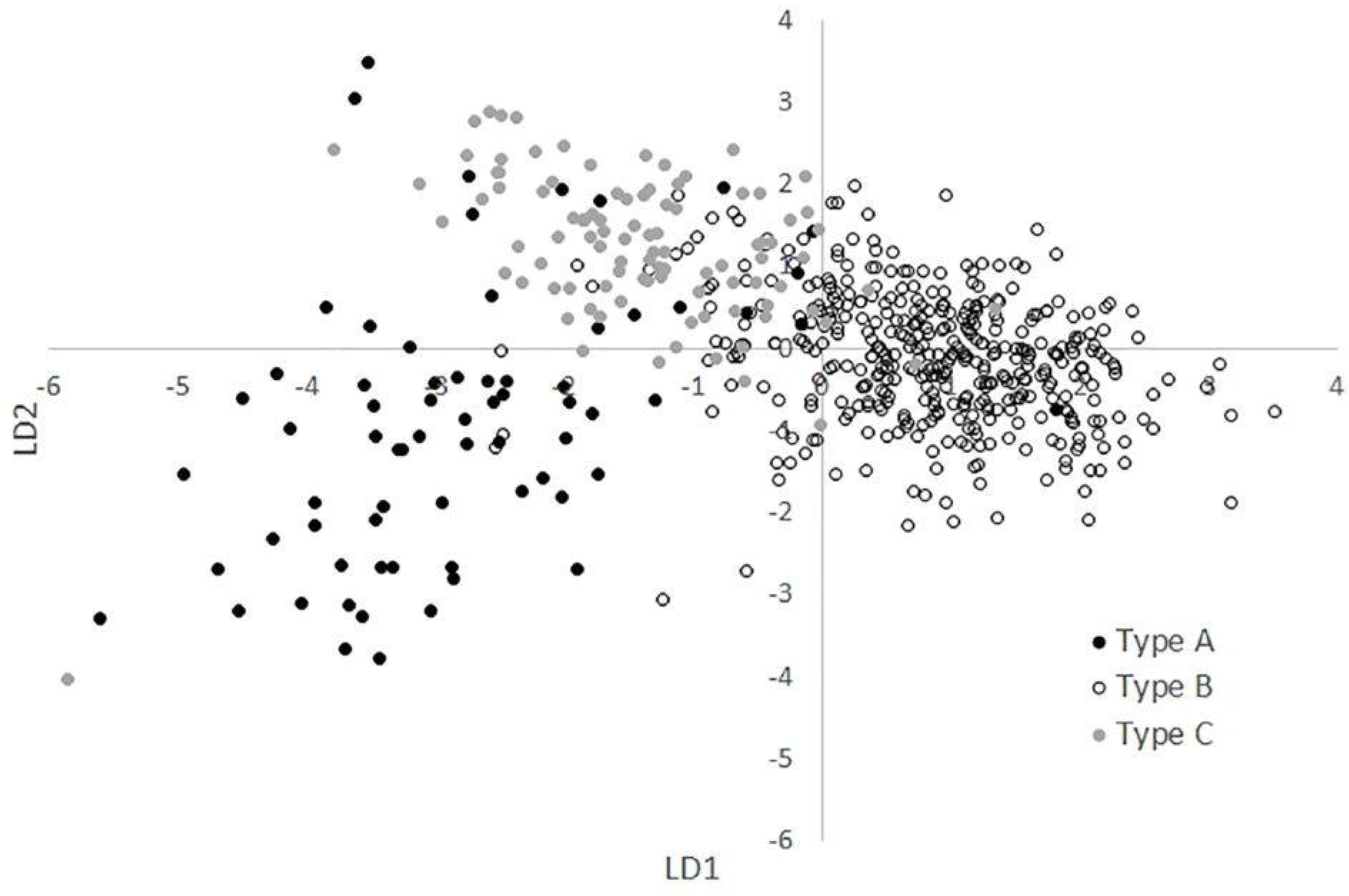
Diagram showing assigned song types (A, B, and C) for Louisiana Waterthrushes 1957-2018.

A comparison with results from a RF analysis confirmed that the two approaches yielded similar results (Table 3). Finally, we plotted misclassifications between the assigned song type and both the LDA classification (Fig 4) and the Discriminant Function (DF) classification (Fig 5) to better indicate where these methods showed disagreement.

**Fig 4.**
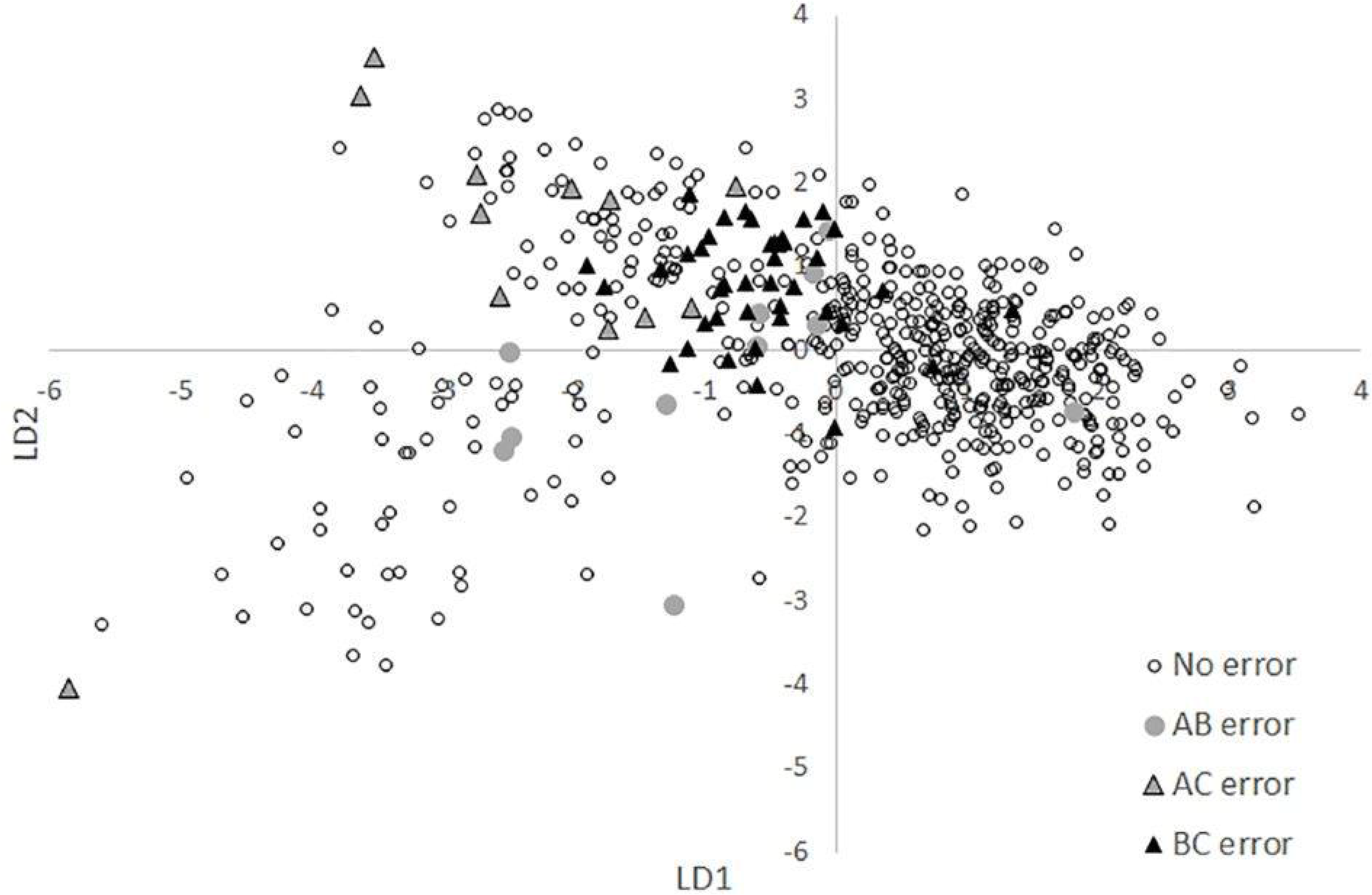
Errors between visual/auditory song type assignments and song type assigned by the Linear Discriminant Analysis. “No error” indicates song types with a match between assigned song type and Linear Discriminant Analysis assignment. “AB error” indicates song types for which the assigned type and Linear Discriminant Analysis type included one Type A and one Type B. “AC error” and “BC error” map errors between those respective pairs of song types.

**Fig 5.**
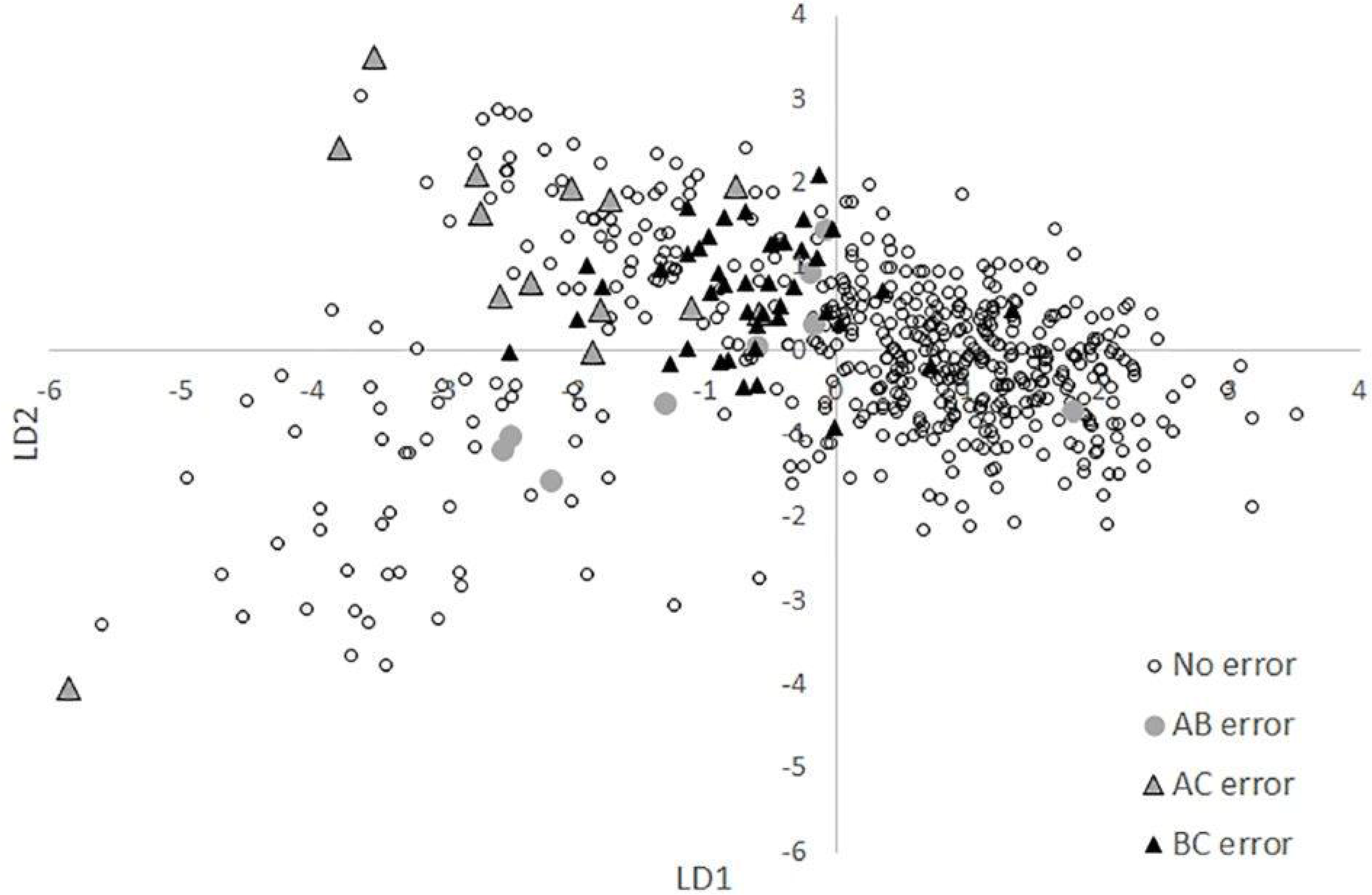
Errors between visual/auditory song type assignments and song type assigned by the Discriminant Function. “No error” indicates song types with a match between assigned song type and Discriminant Function assignment. “AB error” indicates song types for which the assigned type and Discriminant Function type included one Type A and one Type B. “AC error” and “BC error” map errors between those respective pairs of song types.

### Within- and between-song variation

To assess the contribution of within- and between-song variation, we used a dataset consisting of 87 individuals (11 Type A, 52 Type B, 15 Type C, and 9 Unassigned type) and a total of 1583 measured songs. The mean number of measured songs per individual was 18 (SD = 7.8, range was 10 to 32). Most variation was between individuals (Table 4) with all ICC > 0.82. For the duration per note and the three frequency measures there was almost no variation either within or between song types.

**Table 4.** Variance components and Intra-Class Correlation (ICC) for within- and between-song variation in Louisiana Waterthrushes.

| Song attribute | Between individuals | Within individual | ICC |
| --- | --- | --- | --- |
| Notes | 1.030000 | 0.078400 | 0.929 |
| Duration | 0.041700 | 0.008920 | 0.824 |
| Dur_note | 0.001810 | 0.000300 | 0.858 |
| Pct_down | 0.017800 | 0.139000 | 0.928 |
| Min_freq | 0.000014 | 0.009840 | 0.934 |
| Max_freq | 0.000031 | 0.000151 | 0.954 |
| Freq_span | 0.000022 | 0.000219 | 0.909 |

### Unassigned songs

Songs with INS note shapes or patterns that were not readily assignable as Type A, B, or C comprised 11.67% (76) of the total and were designated as Unassigned. The INS of the Unassigned songs sorted into three subcategories: “mixed types”, with notes incorporated from two or in one example three types (38; 5.83%), “equivocal”, which were open to more than one interpretation (30; 4.61%), and “anomalous”, involving note shapes or patterns atypical of Louisiana Waterthrush (8; 1.23%) (Fig 6). The geographic distribution of “mixed types” songs was range-wide, although when displayed on the map concurrently with the A, B, and C types, 30 out of 38 “mixed types” singers occurred in areas where two or three types were present whose INS notes corresponded to notes employed by “mixed types” singers in the vicinity (https://www.google.com/maps/d/u/0/viewer?mid=1PAkxNYqk_7iRslxEd57mgqR38u0&ll=40.669561163969384%2C-83.80830025&z=5). Singers of the undetermined “equivocal” songs had a distribution that conformed with the distributions of Types A and C and were absent in most of the LGM deglaciated zone where Type B predominated. At least half of the “equivocal” songs had INS patterns that included what appeared to be variations on one or more Type A note pairs but these note pairs lacked the symmetry of alternating upstrokes and downstrokes typically seen in the Type A note pair pattern and so were left as Unassigned. The eight “anomalous” singers mostly occurred in the eastern part of the range and consisted of five singers with INS patterns that were a rapid series of abbreviated upstrokes, and three singers that lacked variation between the minimum frequencies of the INS and SNS notes in possible examples of SNS notes substituted for the INS series.

**Fig 6.**
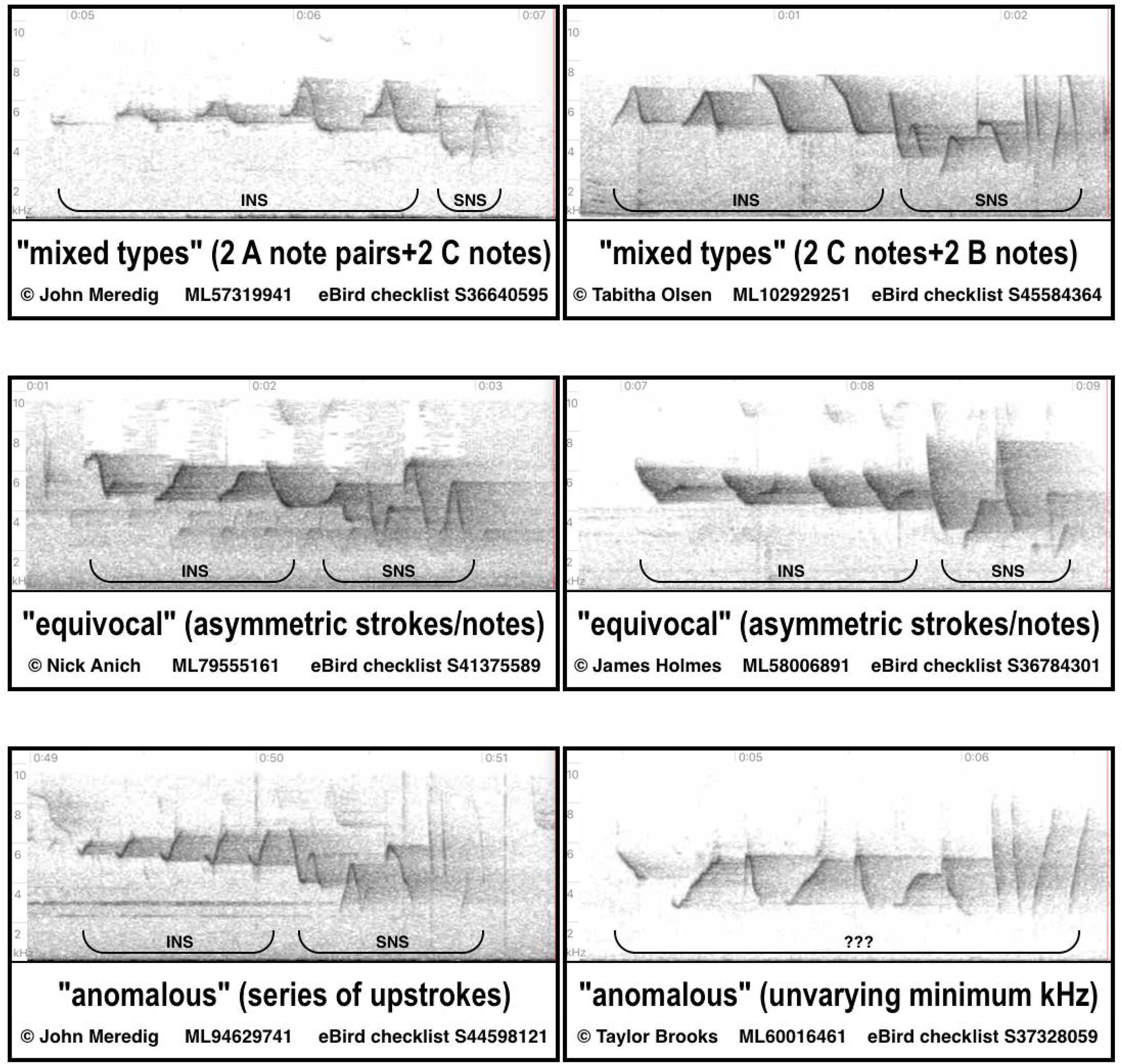
Spectrogram examples of Unassigned songs. Two examples (from S1 Appendix) were selected to represent each of the three subcategories.

### Song type distributions and comparisons with glacial environment

Markers plotted on the map to denote the song locales showed Types A, B, and C had discrete geographic distributions that overlapped in some but not all areas (Fig 7). A depiction of Pleistocene glacial maxima overlaid on the map indicated all three types were distributed widely across previously unglaciated areas of the current breeding range with the exception of the Mississippi Embayment where occurrence was limited to Crowley’s Ridge (Fig 7). The majority of Type B singers (219; 54.2%) were located in the deglaciated zone of the LGM where that type was exclusively ubiquitous. Conversely, occurrences in the LGM deglaciated zone were rare for Type A (2; 2.8%) and minimal for Type C (20; 20.2%) with most of those occurrences recorded near the East Coast (Table 5, Fig 7). Type A singers had notable populations in the Maryland region and Southern Appalachians and were almost entirely absent between the Appalachians and the Embayment but predominated west of the Embayment in the region that remained ice-free during the LGM. Type A was the only song type found along the southwestern edge of the breeding range with the exception of two Type B singers at the range’s southwest terminus. Type C singers occurred primarily east of the Embayment with some located in the LGM deglaciated zone, but the majority were on the unglaciated side of or abutting the Illinoian glacial maximum boundary, and this song type was the only type that occurred directly northeast of the Embayment. A few Type C singers were also observed west of the Embayment where their limited distribution more closely paralleled the distribution of Type A than Type B.

**Table 5.** Percentage of each song type distributed north and south of the Last Glacial Maximum boundary.

| Song Type | North | % | South | % |
| --- | --- | --- | --- | --- |
| A (N= 72) | 2 | 2.8 | 70 | 97.2 |
| B (N= 404) | 219 | 54.2 | 185 | 45.8 |
| C (N= 99) | 20 | 20.2 | 79 | 79.8 |

**Fig 7.**
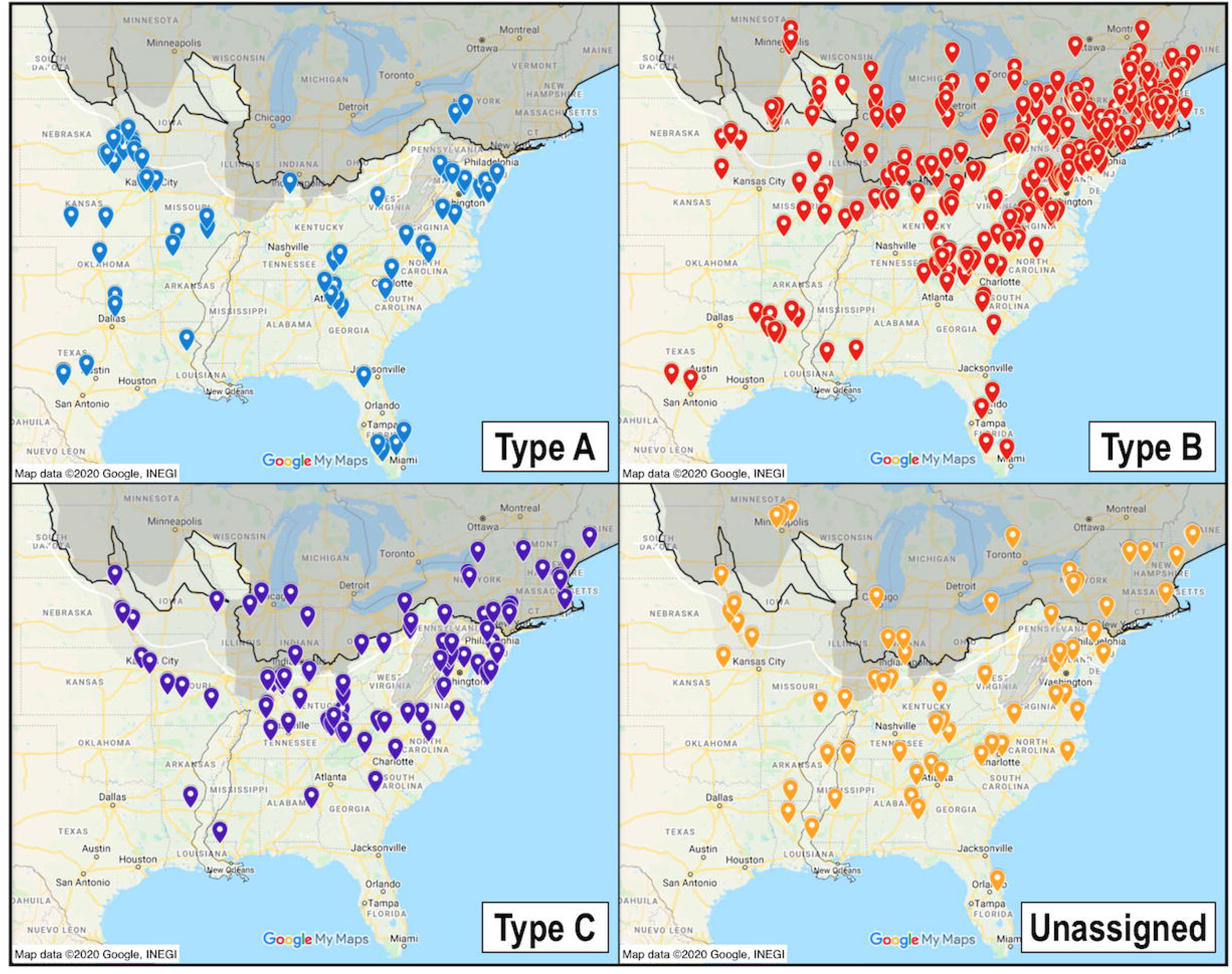
Mapped song type locations with glacial environment. Distributions of the four designated categories relative to pre-Illinoian and Illinoian glacial maxima and high-altitude periglacial permafrost represented by gray screens, and to Last Glacial Maximum boundary indicated by a thick black line, with thin black outline representation of the Mississippi Embayment.

## Discussion

The INS portion of most Louisiana Waterthrush songs consisted of a repetitive series of one of three note shape types that were assigned as Type A, B, or C. Statistical analyses supported assignments obtained from visual and auditory assessment and identified strong predictors of each type. Within the broader structure of each type’s note shapes, there was considerable variance between individual singers in note sizes and dimensions. This individuality within the species’ songs allowed for a multitude of singers to be identified and catalogued from the burgeoning supply of publicly available audio spectrograms, thus offering a largely untapped opportunity to study lineages and dispersal tendencies through unique song variations that are presumably culturally transmitted between generations.

Some insight into the assembly of Louisiana Waterthrush song was provided by examining the INS of “mixed types” singers in the Unassigned category. Twice as many “mixed types” singers sang a combination of notes from Types B and C as singers who combined Types A and B or Types A and C, the reason for which is unknown, but one factor may be Type A’s near absence in the deglaciated zone where Types B and C co-occur (Fig 7; https://www.google.com/maps/d/u/0/viewer?mid=1PAkxNYqk_7iRslxEd57mgqR38u0&ll=40.669561163969384%2C-83.80830025&z=5). Each “mixed types” singer delivered an individually unique combination and pattern of INS notes with a level of consistency between repetitions on par with singers of Types A, B, and C songs. It is informative that most were recorded in the vicinity of two or three neighboring song types whose INS notes corresponded to INS notes in the “mixed types” songs nearby. The parsimonious explanation for these songs with a mixture of types in the INS is a scenario in which juveniles were exposed to adult males singing different song types on adjacent territories during a crucial learning period. In personal observations (S1 Appendix), adjacent territories were most frequently defended by males that shared the same INS type, but it was not uncommon for males of differing INS types to also vigorously defend adjacent territories. The low percentage (5.83%) of total songs analyzed that have a mixture of types in the INS is perhaps unexpected given the prevalent sympatry of the three discrete song types. Careful inspection of the audio spectrograms indicated several instances of counter singing males with differing song types but the presence of more than one type on a stream typically did not result in any known neighboring “mixed types” singers. This observation leads to questions relating to song convergence: why is there a paucity of songs with INS patterns comprised of more than a single type, and why hasn’t exposure to the different types sung by neighbors over the course of time resulted in more Louisiana Waterthrush singers having incorporated various combinations of INS note types as the “mixed types” singers have done? A possible explanation for range-wide maintenance of distinct INS patterns consisting of a single type might be female preference for uniform rather than “mixed types” INS patterns. This plausibly could be similar to dialect recognition in female White-throated Sparrows or what is seen between the taxonomic pairs of eastern and western Marsh Wrens and Eastern and Western meadowlarks when the sympatric occurrence of those phenotypically similar congeners results in females reserving receptivity for singers of their parental song type while singing males evoke territorial defense from both taxa despite their contrasting songs that perhaps could be analogous to contrasting INS patterns in Louisiana Waterthrush. Regarding female Louisiana Waterthrush INS discrimination, we note here an observation of a “mixed types” singer that was defending a territory on which juveniles were present, indicative of a “mixed types” singer having successfully attracted a female who may or may not have shared the same “mixed types” lineage. Limited habitat at that locale accommodated only one male rival who sang a corresponding “mixed types” song; this finding of adjacent singers that shared a unique “mixed types” song pattern suggests a male had returned to his natal stream to vie for territory with a related male. While some shared song patterns (INS + SNS) within the song types were found to occur a considerable distance apart, with the greatest distance noted in a shared Type B pattern found in the deglaciated zone 906 km (563 mi) apart, shared song patterns of all the types were also commonly found on adjacent territories or streams, suggesting that as with the matching “mixed types” neighbors, dispersal may be limited at times, a trait known to contribute to geographic discontinuities in song structure.

The widespread and often overlapping geographic distributions of three statistically separable variations in INS type raise other intriguing questions such as what factors could have contributed to the distributional differences between them? Latitudinal differences place Type B’s distribution northernmost, Type A’s distribution extending northwards the least, and Type C’s as intermediate between Types A and B. Although the three distributions seem indistinctly discrete when cursorily examined, the addition of glacial maxima boundaries to the map in conjunction with the song type distributions reveals possible correlations that may offer clues to each type’s evolutionary history by shedding light on previous colonization opportunities during different glacial and interglacial periods (Fig 7). Type B has a large numerical majority in the area most recently covered by glacial ice and also appears to have the greatest presence in the formerly uninhabitable region of high-altitude periglacial permafrost in the Appalachian Mountains. It was pointed out by Li et al. [77] that postglacial migration of species almost certainly involved more than simple range expansion from south to north, with one scenario having involved expansion from northerly glacial refugia; the current distribution of Type B singers may be illustrative of such alternative expansions considering that type’s current predominance on what would have been newly available habitat following the recession of the LGM. Type A may have undergone a similar expansion during an earlier interglacial period in the same region as Type B’s current distribution to the north of Type C’s geographic stronghold, and this proposition could provide an explanation for a puzzling gap in Type A’s distribution between the Appalachians and the region west of the Mississippi Embayment (Fig 7). Type A’s preponderance on the west edge of the breeding range could plausibly be explained if a succeeding glaciation consistent with the Illinoian glacial maximum had then isolated a subpopulation during convergence of the ice sheet with the Embayment, and facilitated an expansion of Type A in the western region at a time when forest habitat would have prevailed in parts of the Great Plains. West of the Embayment’s bisection of the distributions, the comparatively small Type A subset is disproportionately represented on previously unglaciated terrain and in the area that has not been glaciated since pre-Illinoian episodes, yet it is nearly absent from the neighboring LGM deglaciated zone populated by Type B (Fig 7). The literature has no indication of heterogeneity in the specialized riparian breeding habitat utilized by Louisiana Waterthrushes and so it is unlikely the discernible demarcation between Type A’s occurrence in areas that have not been glaciated since pre-Illinoian episodes and Type B’s major presence in the adjacent LGM deglaciated region can be attributed to differences in the present-day habitat of those contiguous parts of the range. Alternatively, a heritable adaptation to past glacial conditions akin to what is seen in Swainson’s Thrush could feasibly account for the difference in the distributions if such an adaptation is exhibited by Type A but not shared by Type B. This rationale for Type A’s distributional congruency with LGM glacial conditions west of the Embayment could also explain its occurrence entirely on previously unglaciated terrain between the east side of the Embayment and the Appalachians, and similarly offer an explanation for the abundance (35) of Type C singers in that same region which occur on the previously unglaciated side of and abut the Illinoian glacial maximum boundary line, with only a few (5) observed in the deglaciated zone across that same longitudinal span (Fig 7). Although Type C’s distribution largely parallels the distribution of Type A in that large majorities of each occur in the unglaciated region or the region that has not been glaciated since the earlier pre-Illinoian episodes, Type C is not as consistent as Type A in its distributional mirroring of glacial maxima, and a portion (20.2%) of Type C singers also occurs alongside Type B singers in the LGM deglaciated zone. Half of Type C’s occurrences in the LGM deglaciated zone were recorded within 136 km (85 mi) of the East Coast but the reason for that greater density within close proximity to the coast, with the remainder of occurrences more sparsely distributed across the deglaciated zone, is not understood. Type B’s overlapping distribution in the deglaciated zone shows a comparable increase in density near the coast.

Geographic distribution is not the only attribute in which Type C displays overlap with Types A and B. Of the three song types, Type A’s INS frequencies extend the lowest on the kHz spectrum and consist of the lowest average minimum and maximum frequencies, Type B’s INS frequencies extend the highest on the song type spectrum and present the highest minimum and maximum average frequencies, and Type C’s INS minimum and maximum frequency measures are intermediate between the song types (Table 1). Although the three INS types have similar frequency spans, their separation in minimum and maximum frequencies and large-scale distributions may bear some resemblance to the large-scale geographic variations in frequency peaks and span of Chowchilla songs that resulted from isolation in Pleistocene refugia. Origins of the INS variations in Louisiana Waterthrush are obscure. The INS patterns of the three types are not restricted to localized occurrence and therefore results of this study do not readily support pattern variance specifically in the INS portion of the song having arisen on non-contiguous locally isolated streams. We were unable to detect any clinal patterns in the INS that might indicate a drift-like process had led to the three variations. The aforementioned merger of the glacial maximum with the Embayment would have provided a southeastern refugial isolation event in which a subpopulation’s INS pattern could have potentially diverged, but it was not the only Pleistocene isolating scenario in that region. Another possibility might be derived from the numerous studies in recent years [41] that have highlighted the potential role of isolation in other glacial refugia as an explanation for current distributions of many organisms, including *Parulidae*, and this explanation could apply to Louisiana Waterthrush if a subpopulation were to have undergone such an isolation period in a northerly refugium in what would have been theoretically favorable circumstances for INS variation to arise. Genetic research in the eastern United States within Louisiana Waterthrush’s breeding range has established that northerly refugia were maintained in the northern Appalachians region and in the Midwest in areas that gave rise to variations in multiple taxa and harbored deciduous trees that could have potentially afforded suitable habitat. A Pleistocene refugium has been indicated for the sugar maple tree species *Acer saccharum* in the Northeast and a subsequent postglacial migration route from this refugium proposed from the periglacial areas of the Pennsylvania region [78] near a congregation of Type A and C singers. Another refugium in the Midwest, where Type B currently predominates, sustained an unglaciated keyhole of habitat in the Driftless Area where several species of mammals, deciduous plants, and amphibians survived [79—83]. The woody vine *Smilax* sp. survived the LGM in a northern refugium in that area, resulting in an isolated population in the Driftless Area separated from other populations east of the Embayment [77].

After evaluating the mapped distributions of the three identified Louisiana Waterthrush song types, we hypothesize that geographic relationships between INS variations and former Pleistocene glacial boundaries in separate parts of the breeding range could represent contemporary patterns in the type distributions that are indicative of separate evolutionary lineages in three subpopulations shaped either wholly or in part by glaciation-related isolation events. Although this hypothesis may offer a reasonable explanation for the differing distributions, support is inconclusive. We hope this initial exploration of the topic will spur further investigation into potential geographic congruency between the INS portion of the songs and former glacial limits, and also into whether the distributions of the SNS portion of the songs represent smaller-scale dialects.

## Acknowledgements

This endeavor would not have been possible without open access to the large number of spectrograms available through the archives provided by The Cornell Laboratory of Ornithology’s eBird and Macaulay Library, and xeno-canto. We thank all of the recordists who unselfishly placed their hard-won spectrograms into the public domain. We also thank Philip Dixon and Tyler M. Harms at Iowa State University for statistical assistance, and we are grateful to reviewers who provided valuable guidance that significantly improved our approach in this manuscript.

